# Caudata macrogenetics: Geographic attributes and lineage age predict global patterns of mitochondrial genetic variation in salamanders

**DOI:** 10.1101/2025.01.24.634750

**Authors:** Luis Amador, Daniele L.F. Wiley, Irvin Arroyo-Torres, Chris X. McDaniels, Esteban O. Rosario-Sanchez, Hannah Farmer, Hannah Bradley, James Erdmann, Tara A Pelletier, Lisa N Barrow

## Abstract

**Aim:** Genetic diversity contains valuable information about ecological and evolutionary aspects of species. Intraspecific genetic variation is shaped by species’ natural history traits and by characteristics of geography and climate within their ranges. Amphibians are of ecological and conservation interest because of their global distribution, deep history, trait diversity, and roles within ecological communities. Here, we studied genetic variation within salamanders to investigate predictors of nucleotide diversity and spatial patterns of genetic differentiation.

**Location:** Global.

**Time Period:** Present.

**Major Taxa Studied:** Salamanders.

**Methods:** We repurposed mitochondrial DNA sequences and ecological data from open-access databases for 220 salamander species. We calculated nucleotide diversity (π) and tested for isolation by distance (IBD) and isolation by environment (IBE). We analyzed these three variables with random forest and phylogenetic comparative methods using 28 predictors expected to be associated with genetic variation.

**Results:** We recovered 8,108 Cytb sequences with their associated geographic coordinates, of which 7,007 sequences were manually curated by us. Range size, lineage age, and sample size were important predictors of genetic variation. We found higher diversity in regions including the Neotropics and central-eastern Europe. The absence of phylogenetic signal in π, IBD, and IBE suggests that genetic variation is shaped by local ecological and geographical factors rather than by shared ancestry.

**Main Conclusions:** Our finding of range size as an important predictor aligns with theoretical expectations that species with larger ranges tend to harbor more genetic diversity. Furthermore, lineage age being an important predictor is in line with the clade-age hypothesis, in which species with longer divergence times have higher genetic diversity because they have had more time to accumulate genetic variation. Our results underscore the importance of integrating spatial data into macrogenetic studies, providing valuable information for future studies and conservation strategies targeting regions with high or low genetic diversity.

## Introduction

Genetic variation plays a foundational role in maintaining biological diversity (Kardos et al., 2021). One significant evolutionary factor that predicts genetic variation within populations is the number of individuals that contribute DNA to the next generation (the effective population size; Wright, 1931). Genetic variation is proportional to population size, as explained by the neutral theory of molecular evolution (Kimura, 1968). Gene flow between populations is another key factor influencing variation because it can add and maintain novel alleles, counteract the effects of genetic drift and selection, and enable population and species persistence (Park et al., 2024). Natural selection also has an important role in affecting neutral diversity of populations through background or linked selection (Corbett-Detig et al., 2015). Understanding this variation and which factors influence it across different geographic and environmental contexts is essential for elucidating population dynamics and will help us identify populations with conservation needs (Pauls et al., 2013). For example, declining levels of genetic variation may be further exacerbated under global change scenarios due to the reduction of suitable habitat for species (Schierenbeck, 2017).

Climatic and geographic factors shape patterns of dispersal and population size through mechanisms such as habitat connectivity, environmental stability, and barriers to gene flow. These processes, in turn, determine the extent and distribution of genetic variation within species (e.g., Hanson et al., 2017). For example, geographic distance can limit gene flow among populations, resulting in patterns of isolation-by-distance (IBD), where genetic dissimilarity increases as populations become more spatially separated (Wright, 1943). Similarly, environmental differences can restrict gene flow by selecting against migrants non-adapted to the new environmental conditions, leading to isolation-by-environment (IBE), which reflects the influence of habitat heterogeneity and ecological gradients on genetic variation (Wang and Bradburd, 2014). Historical climatic fluctuations and complex topography often act together to promote cycles of population contraction and expansion, which generate and maintain genetic variation. In the alpine toad *Scutiger ningshanensis*, for instance, repeated glacial cycles during the Pleistocene, combined with uplifts of mountain ranges, likely fragmented populations and created refugia, increasing opportunities for genetic differentiation (Meng et al., 2014). Life history traits, particularly those related to reproduction, also influence genetic diversity by affecting effective population size and gene flow (Ellegren and Galtier, 2016). For example, species with high fecundity, widespread dispersal, or external fertilization can maintain large and connected populations, which reduces the effects of genetic drift and promotes diversity. Broad-scale analyses across hundreds of plant and animal species further support the idea that reproductive traits and climatic variables jointly shape genetic diversity by influencing demographic stability and dispersal potential (De Kort et al., 2021), suggesting that these variables act as consistent predictors of genetic variation across taxa.

Studies on focal taxa within a limited geographic range provide the resolution needed to understand local genetic patterns and idiosyncrasies (e.g., Fonseca et al., 2021). It is equally important to investigate genetic variation on broader geographic and taxonomic scales, to have a global perspective of the role of ecological and geographic factors on evolutionary patterns.

Macrogenetics is an approach to analyze open-access DNA sequences across large scales and investigate patterns of intraspecific genetic diversity across higher taxonomic groups (Blanchet et al. 2017). Macrogenetic research has gained popularity over the past decade, driven by the increasing availability of molecular resources (e.g., GenBank), computational tools (e.g., machine learning), and trait databases (Leigh et al., 2021). Despite limitations, particularly the genetic markers represented by available data (see Millette et al., 2021; Paz-Vinas et al., 2021), efforts to understand large-scale patterns and predictors of intraspecific genetic diversity have been undertaken across various taxonomic groups. A study of global genetic diversity in insects attributed high levels of mitochondrial DNA (mtDNA) genetic diversity in the subtropics to temperature and climatic stability (French et al., 2023). Another study of nucleotide diversity using mtDNA for animals and chloroplast DNA for plants in over 38,000 species found that latitude significantly correlates with genetic diversity, with tropical species exhibiting higher intraspecific diversity (Fonseca et al., 2023). These studies highlight the potential of macrogenetics to uncover important predictors of intraspecific genetic diversity across broad sets of taxa globally.

Amphibians contain remarkable diversity in life history strategies and the geographic and climatic variability of their habitats. Recent studies of amphibian genetic diversity indicate that biogeographic region can influence which factors predict intraspecific genetic variation. For example, in Nearctic amphibians, taxonomic family, the number of sequences, and latitude were key predictors of intraspecific variation (Barrow et al., 2021), whereas range size, elevation, latitude, and precipitation predicted genetic variation in Neotropical amphibians (Amador et al., 2024). In the Americas, Lawrence et al. (2023) found a strong relationship between environmental variables and amphibian expected heterozygosity (*H_E_*), and that amphibian *H_E_* decreased with sample size (number of individuals). Within amphibians, salamanders are particularly valuable for macroevolutionary studies because of their broad distribution (excluding Oceania, the Afrotropics, and Antarctica), manageable number of species, and unique traits.

They display remarkable ecological and life history diversity that can directly shape population dynamics and, consequently, genetic structure. For instance, body size variation, from the tiny *Thorius arboreus* (∼20 mm) to the 1.8-meter-long *Andrias japonicus,* can influence dispersal ability and home range size, thereby affecting gene flow and population structure. Likewise, salamanders include generalist species like the tiger salamander, *Ambystoma tigrinum*, versus habitat specialist species adapted to aquatic, terrestrial, or arboreal environments, which can promote or constrain connectivity across landscapes.

One trait that may influence patterns of genetic diversity in salamanders is developmental mode. Species with direct development (i.e., without a larval stage) often have limited dispersal, which could lead to smaller, isolated populations favoring population differentiation (Paz et al. 2015; Liedtke et al. 2022). In contrast, species with a larval stage and dispersing terrestrial adults may maintain more interconnected populations, reducing genetic differentiation (e.g., Zamudio and Wieczorek 2007). Some salamanders exhibit paedomorphosis, a reproductive strategy in which adults retain larval characteristics, which reduces gene flow by allowing persistent aquatic life cycles in species that could otherwise be terrestrial (Denoël et al., 2005). Recent studies have begun to untangle the relationships between species-level traits and genetic variation. For example, Segovia-Ramírez et al. (2023) used genomic data from 62 Neotropical salamander species (one individual per species) to demonstrate that precipitation variability and snout-vent length (body size) were significant predictors of genomic diversity, perhaps related to their roles in shaping population persistence and abundance. Parsons et al. (2024) combined genetic, geographic, climatic, and life history data from salamanders and used a machine learning model to identify predictors of unrecognized genetic lineages. They found that Caudata hidden diversity is the result of variation in climatic variables. A more comprehensive, georeferenced view of intraspecific genetic variation in salamanders, encompassing more species and potential predictors (e.g., climatic and natural history traits), would provide useful insights into the evolutionary and demographic processes driving variation in this group.

Here, we combine molecular, phenotypic, and environmental open-access data to investigate the determinants of intraspecific genetic variation in salamanders. We used machine learning techniques and phylogenetic comparative methods to determine whether natural history traits or climatic or environmental variables can predict genetic variation within species on a global scale. We hypothesized that reproductive traits, such as the total number of eggs (a proxy for fitness) and reproductive mode (e.g., larval vs. direct development), would predict intraspecific genetic variation in salamanders. Species that produce more offspring may maintain larger, more stable populations, which can preserve higher levels of genetic diversity over time (Lacy, 1987; Reed and Frankham, 2003). Additionally, reproductive mode could influence dispersal, leading to lower expected connectivity and more genetic differentiation in species with direct development compared to those with aquatic larvae. We also predicted that traits associated with total species population size (e.g., range size) would be positively correlated with genetic variation, as larger ranges may support more numerous and genetically diverse populations (Frankham, 1996). We hypothesized that species with older evolutionary age (lineage age) may have higher genetic diversity because they have had more time to accumulate variation than younger species (the clade-age hypothesis; see McPeek and Brown, 2007; Scholl and Wiens, 2016). Finally, we expected high spatial autocorrelation in nucleotide diversity of salamanders overall based on their limited dispersal capabilities, which can lead to genetic clustering at small scales (Fusco et al., 2021).

## Methods

### Data collection

We followed the taxonomic classification proposed by AmphibiaWeb (amphibiaweb.org) and Amphibian Species of the World (https://amphibiansoftheworld.amnh.org/) and prepared a detailed count of all species included in this study (Supplementary Material 1, SM1). These databases were also used to reconcile species names and verify taxonomic consistency across datasets. When discrepancies or outdated names were found, we updated species names to match current taxonomy, with a description of our decisions in SM1. We repurposed Caudata DNA sequences, natural history traits, and geographic and environmental data from open-access databases. Following the approach of recent research on the determinants of genetic diversity of multiple amphibian species (e.g., Barrow et al., 2021; Amador et al., 2024), we assembled sequences of the cytochrome-b (Cytb) mitochondrial gene. This gene is suitable for genetic diversity studies because it contains both conserved and variable regions, is commonly used in amphibian single-locus phylogenetic and population genetic studies, and is therefore the most abundant gene in open-access databases for amphibians. Sequences were obtained from GenBank (National Center for Biotechnology Information, NCBI), phylogatR (Pelletier et al. 2022), the *phruta* R package (Román Palacios, 2023), and Amador et al. (2024). We included species with at least 10 sequences, seeking a better representation of intraspecific genetic variation. Our dataset contained species with high variation in sampling effort, such as those with more than 200 sequences. To account for this sampling bias, we also conducted a set of analyses on randomly subsampled datasets that included 10–20 sequences per species. Alignments were generated, visualized, and edited with the MUSCLE aligner v.3.8.31 (Edgar, 2004) in AliView v.1.28 (Larsson, 2014) and saved in FASTA format.

The geographic coordinates (latitude and longitude in decimal degrees format) previously associated with each Cytb sequence were recovered from GenBank, phylogatR, and Amador et al. (2024); corresponding to 31 species (14.1% of the total species) and 1,169 sequences (14.3% of the total sequences). We then manually retrieved the remaining geographic coordinates associated with the sequences (corresponding to 189 species and 7,007 sequences) following a tutorial prepared for this purpose (see Supplementary Material 2). We georeferenced localities using either the function geocode() of the *tidygeocoder* R package (Cambon et al., 2021) or the GEOLocate web application (Rios and Bart, 2010). We only retained sequences that were georeferenced with an error of less than 20 km, resulting in 12 species with fewer than 10 georeferenced sequences.

We calculated both the total range size and the sampled range size of DNA sequences for each species as follows. For total range size, we obtained the geographic range map in shapefile format for each species from the International Union for Conservation of Nature (IUCN) portal using the getIUCN() function of the *rasterSp* R package (RS-eco, 2023). Species ranges not available in the IUCN portal (24 species) were generated with minimum convex polygons using the sequence coordinates and the function mcp() in the R package *adehabitatHR* (Calenge, 2024; see Supplementary Material). This same approach was also used to obtain the geographic range of the sampled sequences for all species (i.e., using only the sequence occurrences). Range size was calculated from shapefiles using the areaPolygon() function of the *geosphere* R package (Hijmans, 2022). Elevation and latitude (min, max, and mean) were obtained from the retrieved geographic coordinates using the function get_elev_point() of the *elevatr* R package (Hollister et al., 2025) for elevation, and custom R scripts for latitude (see Supplementary Material 3). We also obtained precipitation and temperature data (mean and standard deviation) from WorldClim v2.1 (Fick and Hijmans, 2017) using the function worldclim_global() of the *geodata* R package v0.6-2 (Hijmans et al., 2024).

We used the same natural history and geographic traits as in Amador et al., (2024) with slight additions and omissions, choosing predictors we expected to be related to genetic variation (e.g., body size, reproductive mode, elevation, latitude, species range size, precipitation and temperature; Table S1). We omitted the variable activity because nearly all salamanders (except three of the 220 species) included in our dataset were nocturnal. Eight new predictors were added to the matrix (28 predictors in total), which included sampling effort/coordinates (i.e., number of sequences with associated coordinates), sequence length/coordinates (i.e., number of base pairs of sequences with associated coordinates), snout-vent length in millimeters (SVL), paedomorphism (whether or not juvenile features are retained as an adult), litter size min (minimum number of offspring or eggs per clutch), litter size max (maximum number of offspring or eggs per clutch), latitudinal midpoint (mean of the minimum and maximum latitudinal values), and lineage age (estimated species divergence times obtained from Stewart and Wiens, 2025) (Table S1). We obtained trait information from the open-access database AmphiBIO (Oliveira et al. 2017), species accounts in AmphibiaWeb (AmphibiaWeb, 2024), and relevant literature.

### Measures of genetic variation

We evaluated overall genetic diversity (range-wide nucleotide diversity) and spatial patterns of genetic variation (IBD and IBE) within salamander species. We calculated nucleotide diversity (π) with the function nuc.div() of the *pegas* R package (Paradis, 2010) for all sequences (π*_A_*) and sequences with associated coordinates only (π*_S_*). Genetic distances (*gendist*) for each species were calculated based on the raw difference between sequences, using the dist.dna() function of the *ape* R package (Paradis and Schliep, 2019). We tested IBD and IBE for each salamander species as follows. Geographic distance (*geodist*) was calculated between each pair of sequence coordinates using Euclidean distance with the dist() function of the *stats* package (R Core Team, 2023) in R. Before calculating environmental distances, we used the function scale() in R to standardize variables measured in different units (e.g., temperature – °C or precipitation-mm). Environmental distance (*envdist*) was calculated between each pair of sequence coordinates based on the 19 variables of the WorldClim v2.1 database using the vegdist() function of the *vegan* R package (Oksanen et al., 2024). This function calculates a distance matrix, selecting all environmental variables using Euclidean distances. We performed Multiple Matrix Regression with Randomization (MMRR) analyses for each species (Wang, 2013). MMRR quantifies the relative effects of IBD and IBE, represented as distance matrices (*geodist* and *envdist*, respectively), in explaining genetic variation, represented by genetic distances. We compared the observed correlations between *geodist* vs. *gendist* (IBD) and *envdist* vs. *gendist* (IBE) to the correlations from 1,000 permutations. Species were coded as ‘Yes’ if they did present IBD and IBE or ‘No’ if they did not based on an assessment of the associated p-values from the permutation tests. To account for multiple comparisons, we applied a strict Bonferroni correction (Bonferroni, 1936) to reduce the p-value for assessing significance. We used the output (yes or no IBD; yes or no IBE) as binary response variables for further analyses.

We mapped π*_S_*to visualize the distribution of genetic diversity across the globe and assess differences among biogeographic realms (Nearctic, Neotropic, Oriental, and Palearctic). Briefly, following Amador et al. (2024), we combined sequences within a species based on a maximum distance of 100 km between coordinates, and calculated π*_S_* for each locality (see Supplementary Material 4). We then mapped π*_S_*for four different grid sizes (1–4), which correspond with one to four decimal degree grids, or approximately 110 km^2^ to 440 km^2^. We also visualized the number of sequences per grid cell for the four resolutions. For these visualizations, we used the R packages *ggplot2* (Wickham, 2016), *ggspatial* (Dunnington, 2023), and *sf* (Pebesma, 2018; Pebesma and Bivand, 2023). Finally, we tested for spatial autocorrelation for π and the number of sequences separately with the Moran’s I test using the moran.test() function in the *spdep* R package (Bivand et al., 2013).

To further understand the potential link between historical demography and the patterns of genetic variation we found, we used a common statistic for evaluating population size change, Tajima’s D (Tajima, 1989), using the tajima.test() function of the *pegas* package. This test is compatible with different interpretations related to natural selection versus demographic change, making it difficult to distinguish among hypotheses (Yang, 2014). Therefore, we evaluate these results for comparison rather than include the statistic as a variable in our primary analyses. We followed Fonseca et al. (2023) in assuming that the demographic change interpretation is likely more relevant for global comparisons. Thus, in our study, we interpret Tajima’s D values close to zero as representative of neutrality (no population size change over time), positive values as evidence of recent population contraction, and negative values as recent population expansion.

### Predicting genetic variation using random forests (RF)

We used random forest (RF) regression to predict π and RF classification to predict IBD and IBE based on a set of geographic range characteristics and ecological variables. We built RF models with the function randomForest() in the *randomForest* R package (Liaw & Wiener, 2002), with 10,000 trees and 100 permutations per model. We used the tuneRF() function to obtain the optimal mtry parameter (number of variables that are randomly sampled as candidates at each split) and then ran a new RF regression or classification analysis using the best mtry value. The data was split into a training set (70% of the data) for building models and a testing set (30%) for making predictions. For RF regression, we assessed models using the R-squared (R^2^) statistic, which represents the proportion of the variance explained by the model, with a higher R^2^ value indicating a better fit and more predictive power. The relative importance of the predictor variables for π*_A_* and π*_S_* was evaluated based on the percentage increase in mean squared error (%IncMSE).

In RF classification, in addition to the original models, we also tried upsampling and downsampling to address the large class imbalance within our datasets (yes IBD = 56 species, no IBD = 161 species; yes IBE = 39 species, no IBE = 178 species; based on MMRR results with Bonferroni correction). For downsampling, the majority cases (no IBD/IBE) were randomly subsampled to equal the number of minority cases (yes IBD/IBE). For upsampling, minority cases were randomly duplicated to equal the number of majority cases. We chose the best model based on the lowest out-of-bag (OOB) error rate. To assess overfitting and improve cross-validation in the model, we split our data into a training set and a test set (70% train, 30% test). To evaluate predictions of the RF classification models, we created a confusion matrix with the confusionMatrix() function of the *caret* R package (Kuhn, 2008) and tested whether model accuracy was significantly better than the no information rate (NIR). Finally, we assessed the relative importance of the variables based on the Mean Decrease Accuracy (MDA) for the RF classification models of IBD and IBE.

### Evaluating predictors of genetic variation with phylogenetic comparative methods

For phylogenetic analyses, we downloaded a phylogeny subset from VertLife.org for the species in our dataset, which represents phylogenetic relationships inferred by Jetz and Pyron (2018).

We used a subset of 180 species for these analyses because 40 species from our dataset are not included in the available phylogeny. Using a custom R script (Supplementary Material 3), we edited the name of 12 species in the tree to match the names in the dataset (e.g., some *Speleomantes* and *Hypselotriton* species). To assess the potential effect of phylogenetic history in our models, we tested for phylogenetic signal in π using Pagel’s λ (lambda; Pagel, 1999) and Blomberg’s K (Blomberg et al., 2003) implemented in the *phytools* R package (Revell, 2012) with the function phylosig(). For IBD and IBE, we used the *D* statistic implemented in the package *caper* with the function phylo.d() to assess phylogenetic signal.

We used phylogenetic independent contrasts (PIC; Felsenstein, 1985) to investigate the evolutionary correlation between π and the most important predictors identified in the RF analyses. We used the pic() function of the *ape* R package to compute PIC for both response and explanatory variables. We then fit linear models in R for both the uncorrected and phylogenetically corrected (PIC) response and predictor variables. We also used phylogenetic generalized linear mixed models (PGLMMs) to analyze binary traits as described in Ives and Helmus (2011). PGLMMs allow us to account for phylogenetic covariance while reducing Type I error rates. This method was used to analyze IBD and IBE as binary dependent variables (0 for no IBD/IBE, 1 for yes IBD/IBE) versus the most important predictors based on RF results.

Because of the nature of our data (binary outcomes), we chose binomial as the error family with a logit link function. We compared IBD vs. sampling range size, number of sequences, maximum latitude, and mean precipitation; and IBE vs. sampling range size, number of sequences, mean elevation, and mean latitude. We implemented this analysis using the function pglmm() in the *phyr* R package (Li et al., 2020), including a phylogenetic random effect to account for the non-independence among species due to shared evolutionary history.

## Results

### Data summary

We obtained 12,961 Cytb sequences from 220 species distributed globally, of which 8,937 sequences were retrieved directly from GenBank, 2,349 sequences from phylogatR, 1,657 from the *phruta* package, and 296 from Amador et al. (2024). Of these, we obtained geographic coordinates for 8,108 sequences from 219 species. A single species, *Plethodon sherando*, did not have associated coordinates, therefore was not included in IBD and IBE analyses; and 11 species had between four and nine sequences with coordinates. The final dataset included 220 species in nine families (representing >26% of species and 90% of families globally), three response variables (π, IBD, and IBE), and 28 predictors (Table S2). All sequences associated with geographic coordinates and R scripts used to retrieve sequences and coordinates are available as Supplementary Material 4.

### Patterns of genetic variation in global salamanders

Nucleotide diversity within species ranged from π*_A_* = 0 to π*_A_* = 0.087, with *Pseudoeurycea lineola* exhibiting the highest diversity when all sequences were analyzed. When considering only sequences with associated coordinates, values ranged from π*_S_* = 0 to π*_S_* = 0.089, with *Desmognathus amphileucus* showing the highest diversity. We found that nucleotide diversity for all sequences (π*_A_*) and for the sequences with associated coordinates (π*_S_*) were highly correlated (Figure S1; Pearson’s correlation *r* = 0.95, *t*(218) = 46.18, p-value < 0.001). Based on this validation, we conducted further analyses using sequences with associated coordinates (π*_S_*). *Bolitoglossa yariguiensis* was the only species with a value of π = 0 (Table S2; Figure S2). We found that π and the number of sequences were not randomly distributed across the world map (Figure 1, Figures S3 – S5). For example, regions with high genetic diversity included the Ecuadorian Amazon and central-southern Mexico in the Neotropics; and central-eastern Europe in the Palearctic. Areas with low genetic diversity were identified in northern North America (Nearctic), western Europe, and central-western China in the Oriental realm (Figure 1).

**Figure 1.**
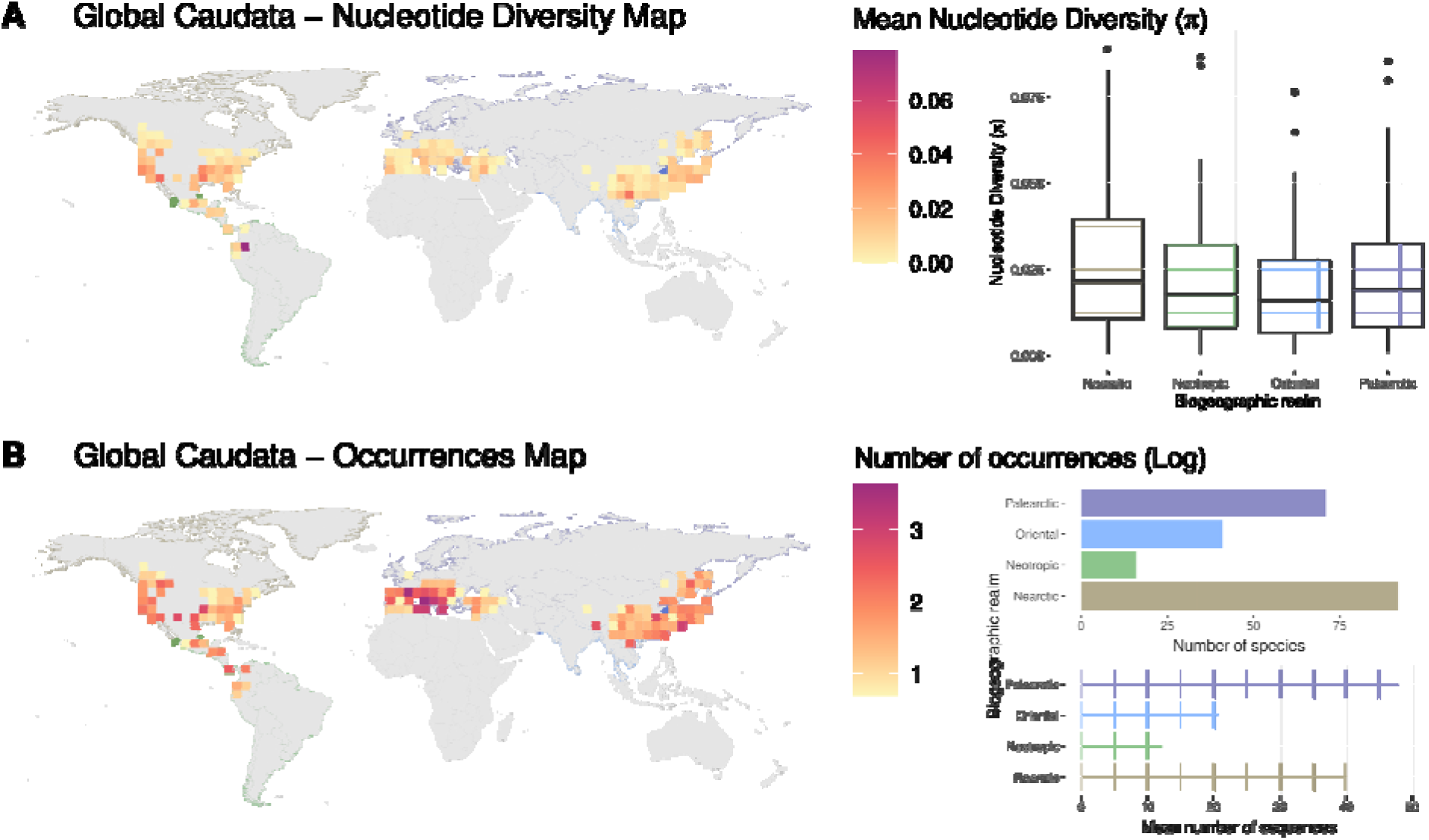
Maps of A) mean nucleotide diversity (π) and B) mean number of sequences with coordinates (log scale) of global salamanders. Biogeographic realms are indicated with shading: Nearctic (brown), Neotropic (green), Oriental (blue), and Palearctic (purple). Boxplots of nucleotide diversity for each biogeographic realm are shown in map A. The number of species and mean number of sequences with coordinates for each biogeographic realm are shown in map B. For visualization purposes, we show grid cell = 4° (approximately 440 km^2^) for mapping.

Furthermore, we found statistically significant, but weak, positive spatial autocorrelation in both π*_S_* (Moran’s I = 0.1435, p-value < 0.0001) and the number of sequences (Moran’s I = 0.4136, p-value < 0.0001).

Most species did not present IBD and IBE patterns, with 161 species (74.2%) showing no IBD and 178 species (82%) showing no IBE (Table S2; Figure S6). We found that only 21 species had significant patterns for both IBD and IBE (Table S2). The Neotropical and Oriental realms had the lowest percentages of species that showed IBD and IBE (IBD: 13.3% and 24.4%, IBE: 13.3% and 10.8% respectively; Figure S7). The Palearctic region had the highest percentage of species that showed IBD (31%) and IBE (25.4%) patterns (Figure S7).

Genetic variation was randomly distributed across the phylogeny (Figure 2). For example, we found species in the same genus with dissimilar π values, such as *Plethodon* (range = 0.0008 – 0.0698) and *Eurycea* (range = 0.0015 – 0.0768) in Plethodontidae, *Paramesotriton* (range = 0.001 – 0.0767) in Salamandridae, and *Hynobius* (range = 0.0008 – 0.0853) in Hynobiidae. We found no or low phylogenetic signal in π based on the method used, for example, with Pagel’s λ = 0.0005, p-value (based on LR test) > 0.05; and Blomberg’s K = 0.053, p-value (based on 1000 randomizations) > 0.05 (Figure S8). We found no significant phylogenetic signal in IBD (Estimated D_IBD_ = 0.945, p-value = 0.249), and significant but moderate phylogenetic signal in IBE (Estimated D_IBE_ = 0.782, p-value = 0.012) (Figure S9).

**Figure 2.**
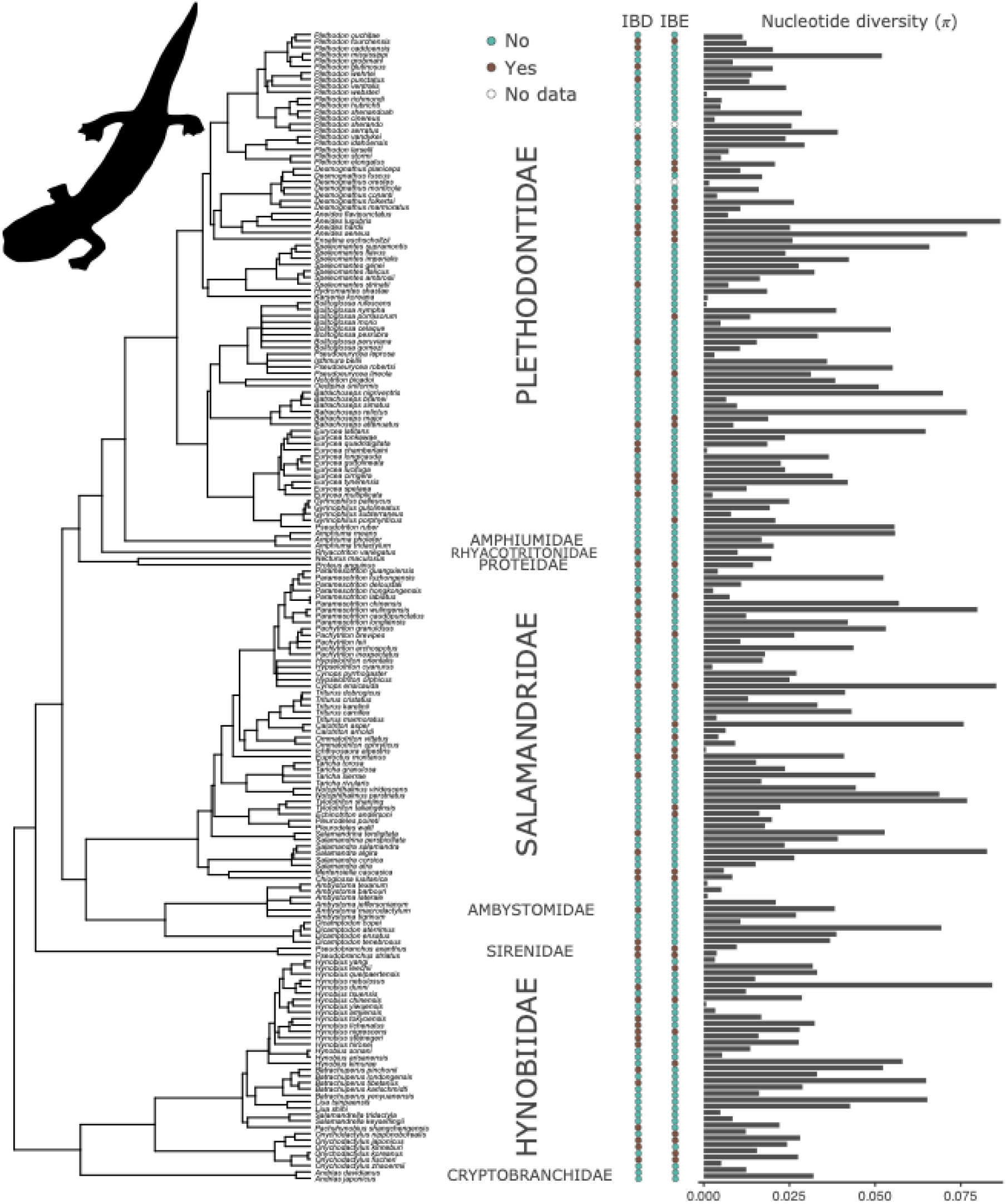
Phylogeny of Caudata species included in this study, showing the values of nucleotide diversity (π) as gray barplots to the right and the presence (brown) or absence (teal) of isolation-by-distance (IBD) and isolation-by-environment (IBE) per species. Relationships are based on Jetz et al. (2018). *Notophthalmus viridescens* silhouette was taken from phylopic.org.

Our results from assessing Tajima’s D indicate that many species in our dataset (n = 109) have undergone recent population expansion (i.e., negative Tajima’s D, more recent mutations than expected under neutrality; Figure S10). Positive Tajima’s D values indicate more alleles at intermediate frequencies than expected under neutrality, possibly because of a recent population contraction or bottleneck, as we observed in 51 salamander species (Figure S10). Fifty-nine species do not show strong evidence for population expansion or contraction (Tajima’s D ≈ 0) suggesting no deviation from expectations under neutrality (Table S2; Figure S10).

### Predictors of genetic variation in global salamanders

Our random forest analyses showed an influence of sampling effort on genetic variation estimates, with the number of sequences (with associated geographic coordinates) as a top predictor of IBD, and IBE (Figure 3). Additionally, we compared both species range size variables in our dataset, total range size versus sampling range size, and we found a strong, positive linear relationship (Figure S11; Pearson’s correlation *r* = 0.87, *t*(218) = 25.82, p-value < 0.001). These results suggest the sampled range for most species adequately represented the total range size. Nevertheless, we ran RF analyses using both total range size or sampling range size and found similar results between the two variables (Figures 3, S12). Because the range size variables were highly correlated with one another and captured similar aspects of the species’ biology, we subsequently describe the results using the sampling range size.

**Figure 3.**
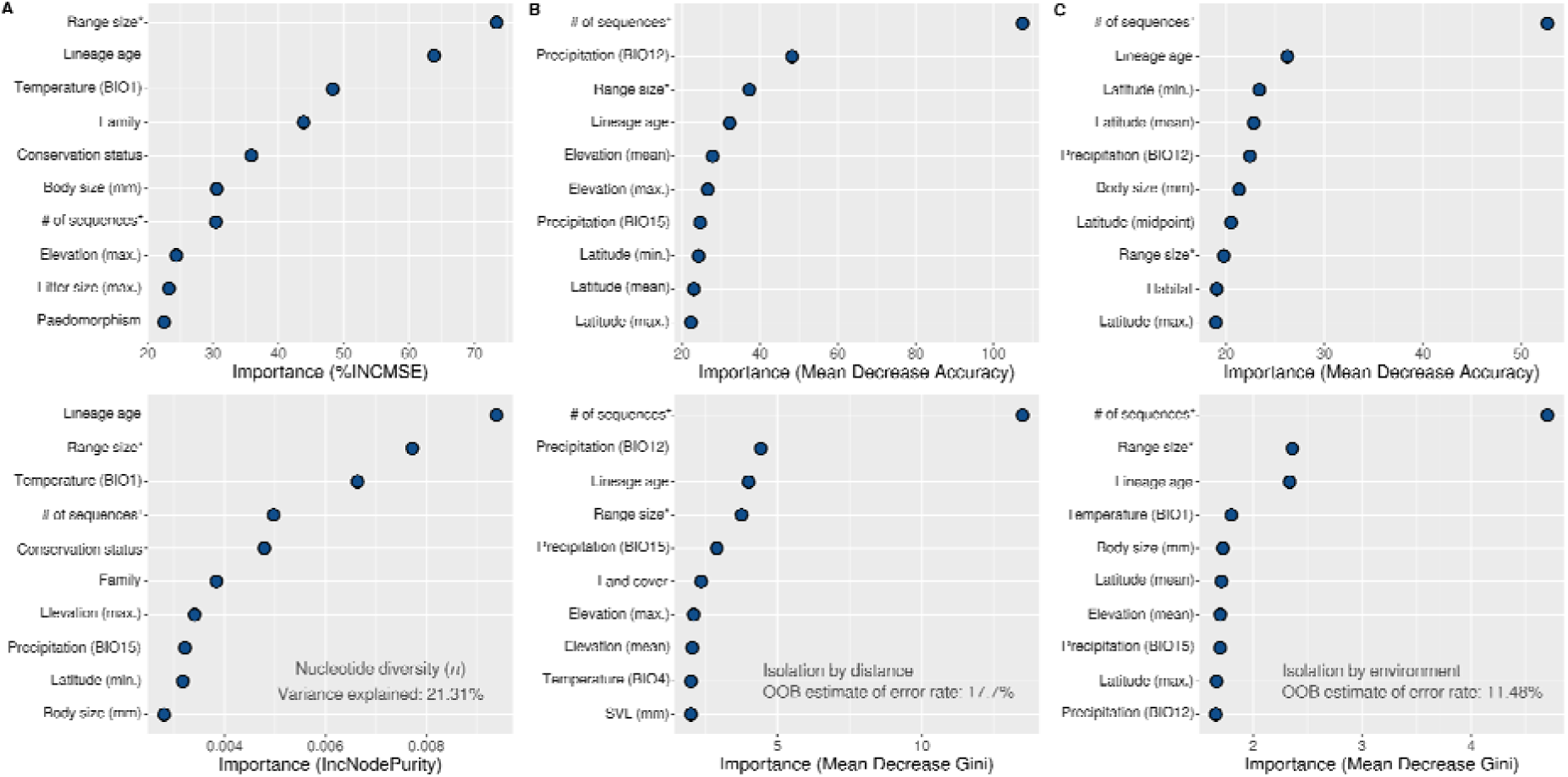
Random Forest (RF) results for A) nucleotide diversity (π, using RF regression), B) isolation-by-distance (IBD, using RF classification), and C) isolation-by-environment (IBE, using RF classification). All predictors except the number of base pairs, and the total range size were used in these RF models, but only the top 10 best predictors for each model are shown here. Range size* = sampling range size based on localities with associated DNA sequences; # of sequences^+^ = number of sequences associated with geographic coordinates; SVL = snout-vent length; mm = millimeters; BIO1 = Annual Mean Temperature; BIO4 = Temperature Seasonality (standard deviation ×100); BIO12 = Annual Precipitation; BIO15 = Precipitation Seasonality (Coefficient of variation).

We found that sampling range size and lineage age were the best predictors of π and were two of the best predictors of IBD and IBE in random forest analyses (Figure 3). Random forest regressions showed that taxonomic family was the fourth-best predictor of π following mean temperature (BIO1) considering the Increase in Mean Squared Error (%IncMSE) measure (Figure 3a, S13). Validation of the RF regression results for π based on the R^2^ score indicated good model performance (R^2^ = 0.95). Mean precipitation (BIO12) and lineage age were other important predictors of IBD (Figure 3b, S14). For IBE, RF showed that latitude was another important predictor (Figure 3c, S15). In RF classification analyses, the best IBD and IBE models were obtained when we downsampled our dataset. Our IBD RF model had high accuracy (88.1%; 95% Confidence Interval CI: 77.82% – 94.7%), low sensitivity (56.25%), and very high specificity (98.04%); our model was statistically better than predicting only the majority class, p-value [Acc > NIR] = 0.0116. The IBE RF model had high accuracy (94.74%; CI: 85.38% – 98.9%), low sensitivity (57.14%), and very high specificity (100%); however, the model’s accuracy was not statistically better than the NIR, p-value [Acc > NIR] = 0.0688.

Sampling range size had a significant, positive relationship with π, showing that genetic diversity within species increases with the geographic area sampled (Figure 4A; R^2^ = 0.09; p-value < 0.001). We found a similar pattern using phylogenetic independent contrasts (PIC), where sampling range size and π had a strong, positive relationship (Figure 4B; R^2^ = 0.11; p-value < 0.001). The relationship between lineage age and π was positive and significant, in which older species had higher genetic diversity in comparison with younger ones (Figure S16A; R^2^ = 0.11, p-value < 0.001), but this correlation was not significant taking the phylogeny into account (Figure S16B; R^2^ = 0.004, p-value: 0.405). We found a significant, positive relationship between mean temperature (BIO1) and π, suggesting that species living in areas with higher temperatures have more genetic diversity (Figure S17A). However, PIC did not show a significant correlation between temperature (BIO1) and π after correcting for phylogenetic relatedness (Figures S17B).

**Figure 4.**
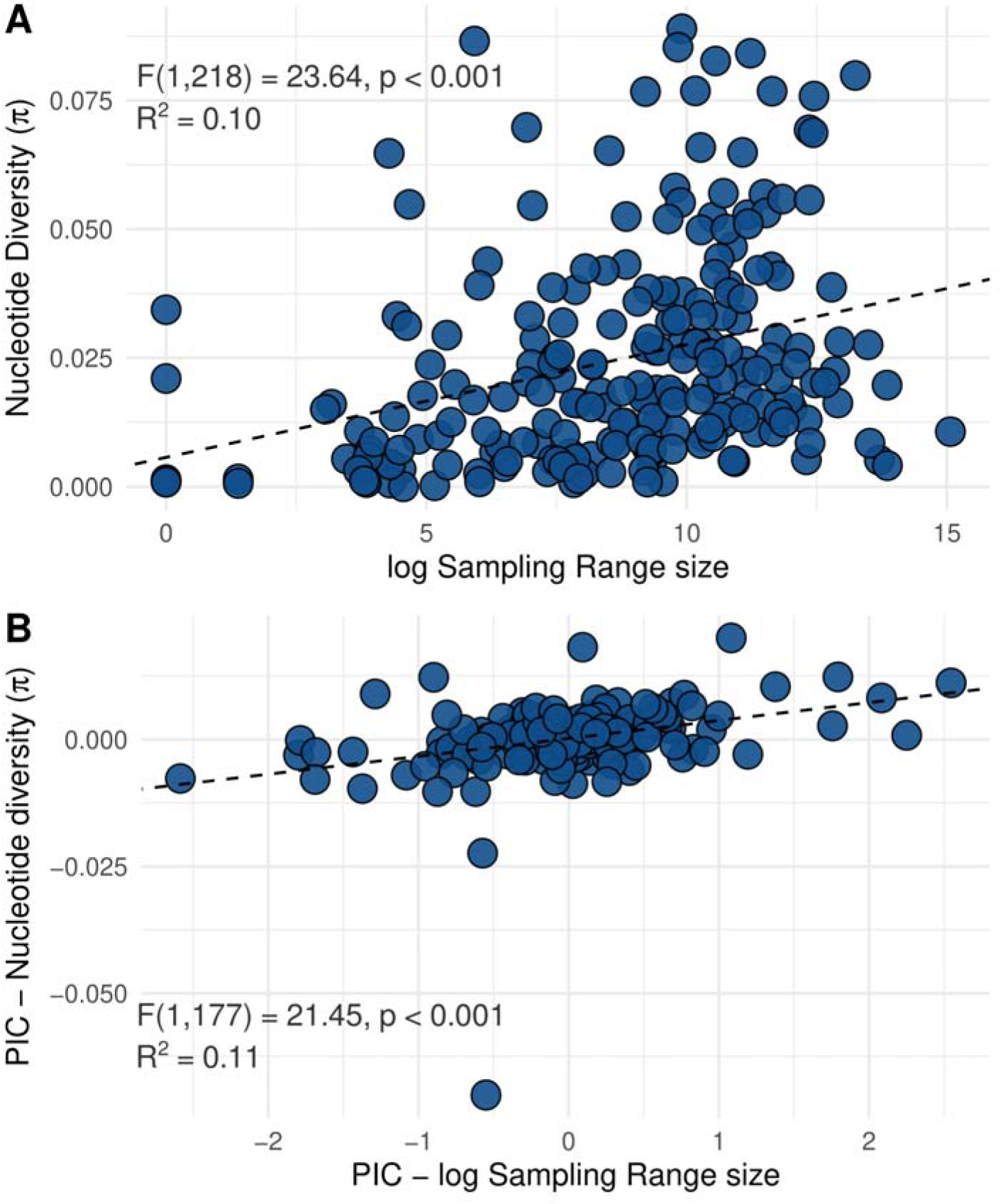
A) Comparison of sampling range size (in log scale) with nucleotide diversity (π) for each species. B) Comparison of phylogenetic independent contrast (PIC) results between the sampling range size (log scale) and nucleotide diversity (π). Linear regression results are indicated in each panel.

The number of sequences associated with geographic coordinates, lineage age, and sampling range size were top predictors for IBD and IBE models. There were clear differences between the number of sequences sampled and IBD/IBE patterns (p-value < 0.01), in which species with more sequences tended to show significant IBD/IBE patterns (Figure 5A, D). Species with significant IBD/IBE had slightly larger sampling range sizes on average (Figure 5C, F) and also tended to have older lineage ages (Figure 5E, S18). When temperate and tropical species were considered separately, we found a similar trend in which species with larger ranges were more likely to exhibit IBD and IBE, but results were not significant (Figure S19). Phylogenetic generalized linear mixed models (PGLMM) found that the number of sequences had a statistically significant, positive relationship with IBD (β = 0.01655, SE = 4.10 x 10 ^3^, p < 0.001), and IBE (β = 0.0158626, SE = 4.27 × 10 ^3^, p < 0.001). These results suggest that as sampling effort increases, the likelihood of detecting an IBD or IBE pattern increases (Table S3). Coefficients, standard errors, Z-scores, and p-values of the other predictors (i.e., sampling range size and precipitation for IBD, and lineage age for IBE) are shown in Table S3.

**Figure 5.**
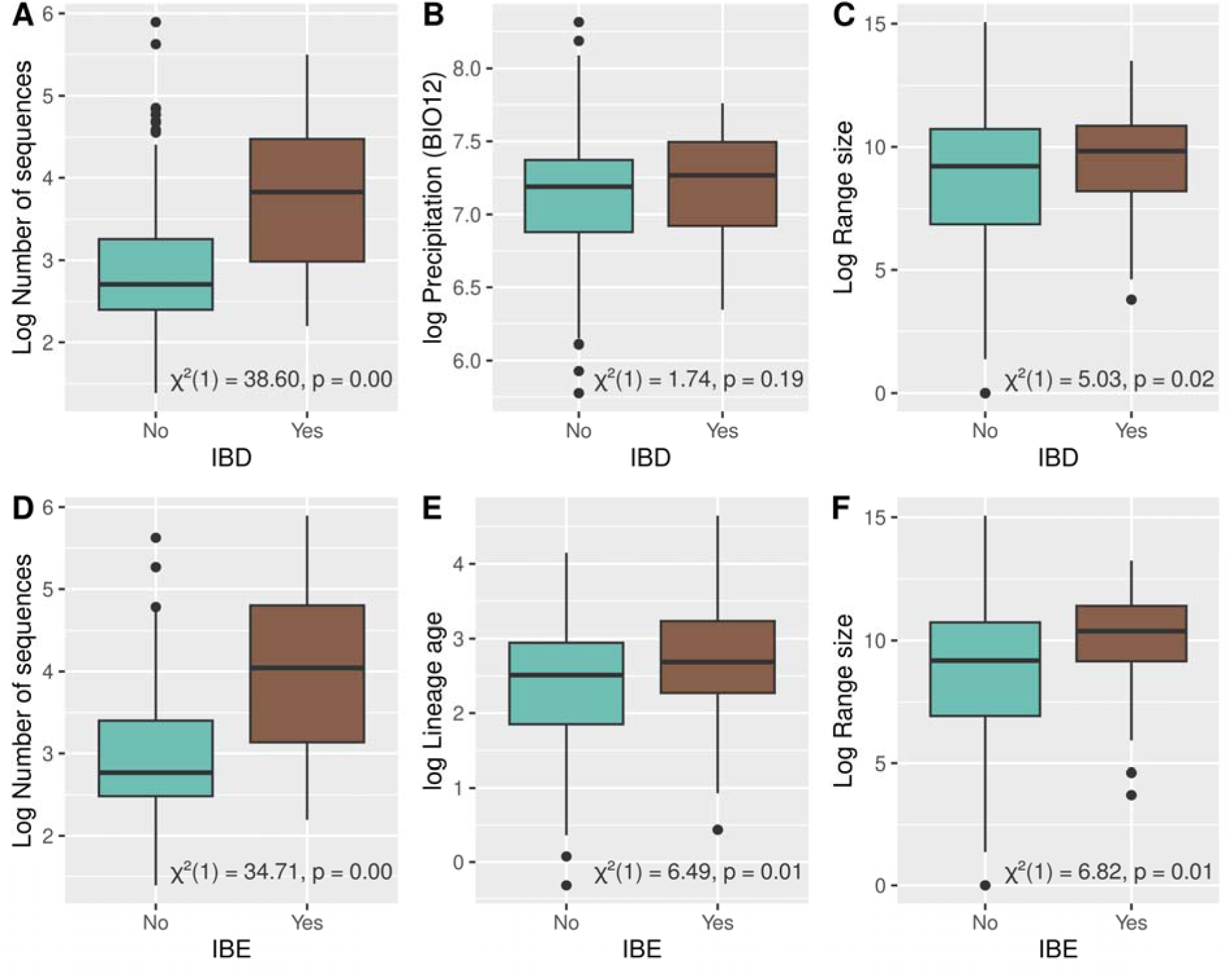
Boxplots comparing the number of sequences (A, D) and the sampling range size (C, F) between species with (Yes, brown color) and without (No, teal color) isolation-by-distance (IBD) and isolation by environment (IBE). Comparisons of precipitation (BIO12) with IBD (B), and lineage age with IBE (E) are also shown. Each panel includes the results of a chi-squared test. Black dots represent outliers.

## Discussion

In this study, we analyzed 12,961 Cytb sequences from 220 global salamander species and identified key predictors of intraspecific diversity (π) and patterns of spatial genetic variation (IBD and IBE). Species range size and lineage age were important predictors of all three aspects of genetic variation. Specifically, species occupying larger geographic areas tended to have greater overall nucleotide diversity and to be more geographically and environmentally structured. The results for π are consistent with population genetics theory (Wright, 1931), in that large geographic ranges can support larger populations that harbor more genetic diversity because of reduced genetic drift (Lacy, 1987). The significant relationship found between range size and spatial genetic differentiation (i.e., IBD and IBE) could be explained by the greater potential for geographic and environmental isolation between populations at distant locations in species with large ranges (Eckert et al., 2008). However, species range size alone does not explain much of the variation in IBD or IBE presence (small effect sizes recovered in logistic regression models), and the strong influence of the number of sequences sampled highlights one of the major challenges for macrogenetic studies—limited data availability for many species worldwide.

Our results provide only partial support for the clade-age hypothesis (e.g., random forest results). Although lineage age showed a positive and significant relationship with intraspecific genetic variation across global salamanders, there is no association when the phylogeny is included (see PIC and PGLMM results). For example, we tested whether older salamander lineages tend to accumulate more intraspecific genetic diversity than younger ones, the clade-age hypothesis (see McPeek and Brown, 2007), and we found that there is a significant non-phylogenetic correlation, but this relationship disappears once phylogenetic relatedness is taken into account. Similar patterns have been observed in other vertebrate groups, where evolutionary time is recovered as one of the best predictors of genetic diversity (e.g., Theodoridis et al., 2020 in mammals). Our results suggests that the observed effect of lineage age on genetic diversity could be explained by shared evolutionary history rather than independent accumulation of variation through time. The strong phylogenetic signal detected for lineage age (Pagel’s λ = 0.9999, *p* < 0.001) indicates that this trait is almost entirely structured by shared ancestry. This high phylogenetic dependence reinforces that comparisons involving lineage age must account for shared ancestry (PIC or PGLMM models) to avoid inflated associations with other traits, such as π, IBD or IBE (e.g., Lai et al., 2025). Thus, our results indicate that spatial (range size) and environmental factors (precipitation, temperature) may play a stronger role than lineage age in shaping intraspecific genetic variation across Caudata. This suggests that long-term ecological and demographic stability, rather than lineage age, may better explain the maintenance of high intraspecific genetic variation in some salamander lineages (e.g., Pan et al., 2019; Iannella et al., 2025). The recovered relationship between lineage age and intraspecific genetic variation may be confounded by lineage-level traits (e.g., life history traits, species range size) that are conserved phylogenetically. In fact, we found a positive and significant relationship between lineage age and sampling range size, as recently proposed by Alzate et al. (2025).

### Global patterns of nucleotide diversity (π)

Our results provide evidence of the uneven distribution of nucleotide diversity (π*_S_*) across the globe. We observed that certain hotspots of genetic diversity, like the Ecuadorian Amazon and central Europe, southwestern and central North America, and southeast Asia, align with regions of high biodiversity (e.g., Mittermeier et al., 2011). Contrastingly, low genetic diversity in regions such as northern North America or western Europe could reflect recent recolonizations or historical bottlenecks (e.g., Riberon et al., 2002; Bonato et al., 2018; Auteri et al., 2022). Our results only partially contrasted with those of Miraldo et al. (2016), who found that amphibian genetic diversity decreased at higher latitudes. Although our map revealed a broadly similar geographic pattern, latitude itself (mean, maximum, minimum, or midpoint) was not an important predictor of genetic diversity in salamanders. Instead, the predictors that best explained genetic diversity, such as species range size and environmental factors, tended to be associated with regions closer to the equator. The high diversity we found in the Neotropics (16 species) could be explained by a combination of factors including tropical climate stability, high topographic heterogeneity, or long evolutionary history in the Neotropics (Antonelli, 2022).

Another plausible explanation for the observed high levels of π in some regions is that they coincide with glacial refugia, areas where species persisted and maintained high levels of genetic diversity through time (e.g., Ramírez Barahona and Eguiarte, 2013). The Andean foothills were a major glacial refugium, particularly the central Andes in the Neotropical region (e.g., Escobar et al., 2021). In the Nearctic, the Southern Appalachians and coastal areas of the Pacific Northwest are considered potential glacial refugia (e.g., Shafer et al., 2010). In the Palearctic, the Caucasus Mountain range has been identified as a potential refugium during glacial periods (e.g., *Triturus* newts; Wielstra et al., 2013). The Korean Peninsula and the mountainous regions of China are two examples of glacial refugia in the Oriental region (Fu and Wen, 2023). Low levels of π in northern North America are consistent with the impact of historical and demographic processes in the region, such as Pleistocene glaciations and associated range contractions in salamanders (Rovito and Schoville, 2017; López-Delgado and Meirmans, 2022). We showed that species experiencing expansion, based on Tajima’s D, presented low levels of nucleotide diversity (Figure S20). This result is expected when recent expansion from a small population occurred after a bottleneck, leading to a limited gene pool carried by a few founding individuals, such as range expansions after the last glacial maximum (Mayr, 2001). Interestingly, the occupation of refugia by salamander species may have contributed to the extinction of older lineages and fostered more recent diversification events, consistent with patterns observed in the regions mentioned above (see Tingley and Dubey, 2012). We observed that species experiencing demographic expansion often have young evolutionary age (Figure S21), reflecting recent diversification events associated with demographic growth.

### Global patterns of isolation-by-distance (IBD) and environment (IBE)

The limited evidence for both IBD and IBE across species (only 21 species, or ∼10%, showed both patterns) suggests that gene flow in many salamander species may not be constrained by geography or environment. Instead, the genetic variation in these species is shaped more by demographic processes such as bottlenecks or population expansions than current spatial or ecological gradients (e.g., Sexton et al., 2024). For instance, some salamander species may maintain high gene flow that can homogenize genetic variation, reducing patterns of IBD or IBE. In contrast, others with strong philopatry may exhibit spatial genetic structure influenced more by historical processes that disrupt patterns of genetic isolation associated with IBD and IBE (Wake, 2009; Sexton et al., 2014). For example, montane populations isolated in separate refugia in response to climatic changes might differentiate and exhibit highly restricted gene flow even at short distances or among similar environments, as reported in plethodontid salamanders (Kozak and Wiens, 2006; Rovito, 2017).

These results underscore the importance of considering biogeographic and ecological context when evaluating the drivers of intraspecific genetic variation on a global scale. The proportions of species showing IBD and IBE differed slightly across biogeographic realms, with the Palearctic including the most species with these patterns. One example is the fire salamander (*Salamandra salamandra*), one of the most common and widespread salamander species in Europe, which was found to be strongly structured in a landscape genetics analysis (Bani et al., 2015). The Neotropical and Oriental regions had lower proportions of species with IBD or IBE patterns compared to the other regions, but these differences were relatively minor. This contrast between realms suggests that historical biogeography and environmental complexity are key to understanding spatial genetic variation. For example, the incidence of IBD/IBE in the Nearctic and Palearctic could reflect the consequence of glacial cycles (Schmitt, 2007), while the low incidence in tropical regions may reflect more recent colonization dynamics that occurred in the late Pleistocene (Carnaval et al., 2009).

### Predictors of genetic variation in global salamanders

Two of the best predictors of nucleotide diversity in this study, the number of sequences and range size, could both be considered proxies of demographic processes. Species with more available sequences could truly be more abundant in nature, reflecting both a true demographic property as well as sampling bias towards species that are more frequently encountered. Either way, sampling more sequences could provide a greater likelihood of detecting genetic variation because more samples analyzed will capture the full range of genetic diversity present within a species. When we analyzed π, IBD, and IBE using a standardized sampling effort per species (10 to 20 sequences), we note that the number of sequences was not an important predictor in π and IBE based on the random forest models, but was the most important predictor of IBD. Similarly, species sampling range size was not a key predictor for IBD and IBE but was an important predictor for π in this subsampled dataset (Figure S22). Lineage age was one of the best predictors for π and IBE in the subsampled dataset (Figure S22). These results suggest that differences in sampling effort across species can provide challenges for macrogenetic studies and should be considered carefully in future studies.

The positive correlations found between number of sequences, range size, and π in our dataset are consistent with the prediction that total population size (with number of sequences and range size as potential proxies) corresponds with genetic diversity (Figure S23). Species with larger geographic ranges can have larger and more balanced populations, reflecting the relationship between effective population size and genetic variation (Waples, 2022). Geographic range size was also one of the best predictors of IBD and IBE, like previous findings in a wide range of taxonomic groups (Pelletier and Carstens, 2018). A large range can harbor more differentiation, where population subdivision could be influenced by different ecological and evolutionary pressures and undergo contrasting demographic histories (Lowe et al., 2017). The relationships between range size and patterns of genetic variation we identified in global salamanders is partially consistent with previous regional studies, such as Amador et al. (2024), which found a similar pattern in Neotropical amphibians for π, but not for IBD or IBE. Similar to previous meta-analyses of IBD and IBE (Crispo and Hendry, 2005; Jenkins et al. 2010), we found that identifying global predictors of spatial genetic variation is challenging because of the many potential factors that influence the presence and detection of these patterns.

Climatic variables, including different aspects of temperature and precipitation, were among the top ten predictors of genetic variation in RF analyses, suggesting an important role of climate in diversity and population differentiation within salamanders. Mean temperature (BIO1) was the third best predictor for π. The association between higher genetic variation and warmer temperatures is consistent with the evolutionary speed hypothesis (Wright et al., 2010; Gillman and Wright, 2014), where genetic diversity is expected to increase with temperature. Variation in precipitation (BIO12) was the second most important predictor for IBD after the number of sequences. A possible explanation for this association is that current stable climates, such as those in tropical environments, promote spatial genetic variation, thus supporting the climate stability hypothesis (Janzen, 1967; Stevens, 1989). Although the relationships between these climatic variables and the response variables we examined were relatively weak, previous work has also highlighted the role of climate in explaining variation within salamanders. Specifically, phylogeographic structure measured using species delimitation methods was best explained by variation in climate (Parsons et al., 2024). Like other studies on birds (Smith et al., 2017) and amphibians (Barrow et al., 2021), our analyses suggest that life history and ecological traits are less important than geographic or climatic variation for predicting intraspecific diversity in salamanders.

### Phylogenetic, taxonomic, and conservation implications

We did not find phylogenetic signal in π, although taxonomic family was the fourth-best predictor of π in the final Percentage Increase in Mean Squared Error (%IncMSE) RF model, suggesting that some variation can be explained by evolutionary history, but not all. Sample sizes were limited within some families, but diversity estimates in those families were consistently high (e.g., Sirenidae) or low (e.g., Cryptobranchidae), indicating family can be a useful predictor of intraspecific variation. On the other hand, discrepancies in π within genera of family Plethodontidae, such as observed in *Plethodon* or *Eurycea*, emphasize the influence of local ecological and recent demographic processes over shared ancestry for influencing variation within species. Processes leading to homoplasy (e.g., convergence, parallel evolution) dilute phylogenetic signal (Klingenberg and Gidaszewski, 2010). Homoplasy is a common phenomenon in salamander evolution; for example, paedomorphosis has evolved independently several times (Wake, 2009). Likewise, levels of genetic variation may be similar in salamander species that do not share a recent common ancestor (Kozak and Wiens, 2006). Phylogenetic signal in IBD was not statistically significant either, suggesting other factors independent of phylogenetic history influence these patterns. This result was consistent with a previous study on Neotropical amphibians (Amador et al. 2024), which found no phylogenetic signal in IBD or IBE. Significant phylogenetic signal in nucleotide diversity, however, was previously found in amphibian datasets including frogs (Barrow et al. 2021; Amador et al. 2024).

Species with higher π values could contain cryptic diversity and would be useful target species for exploring species boundaries in future studies (e.g., Parsons et al. 2022). Using a smaller dataset than the present study (83 salamander species total, most from the COI mitochondrial gene), Parsons et al. (2024) found that ∼2/3 of species in the dataset showed strong phylogeographic structure, determined by single locus molecular species delimitation methods. Several species with high values of π in our dataset match species from their study, indicating they have hidden genetic lineages that should be further explored (*Mertensiella caucasica, Bolitoglossa rufescens, Plethodon cinereus, Eurycea cirrigera, Batrachoseps attenuatus, Ommatotriton vittatus, Aneides flavipunctatus*), ideally incorporating nuclear genomic data and thorough geographic sampling.

Salamanders are within one of the most threatened vertebrate groups (Luedtke et al., 2023) and require innovative syntheses of available information to highlight priorities for conservation strategies. Nucleotide diversity could be a useful proxy for conservation status (Petit-Marty, 2021). The relationship we identified between species range size and genetic variation highlights the vulnerability of species with small ranges. Conservation status was an important predictor for π, but sample sizes were limited within several categories. Seventy-seven (35%) of the 220 species in our dataset are threatened based on the IUCN Red List; 12 as Critical Endangered (CR), 30 Endangered (EN), and 35 Vulnerable (VU). These salamanders should receive special attention, especially if they occupy small ranges and have low genetic diversity.

### Limitations and challenges

One major limitation for macrogenetic studies is the uneven spatial sampling of available sequences, which can lead to an incomplete picture of the distribution of genetic diversity. There are also sampling biases among taxa, such that some species are well characterized while others are omitted from global comparisons. Another limitation is the use of a single mitochondrial gene to calculate genetic diversity and genetic distances. Although mtDNA is widely used in phylogenetic and population genetic studies because it evolves quickly (Galtier et al., 2009), it provides an incomplete picture of intraspecific diversity because it is a single locus, haploid, and solely maternally inherited (Moore, 1995). Moreover, although there is a potential association between lineage age and genetic diversity, it is important to note that divergence times estimated from different loci can have distinct coalescence histories and are often younger than species ages. This suggests that a relationship between divergence times across loci and genetic variation inferred from a single locus (mtDNA) is possible but not necessarily straightforward (Pie and Caron, 2023). However, single locus macrogenetic studies are still valuable because the amount and availability of data elucidate patterns associated with historical processes and variability within species. Furthermore, the large genome size of salamanders hampers the use of nuclear genomic data in studies on this scale. These limitations reinforce the need for additional geographic sampling of high-quality tissues that include broader taxonomic coverage and facilitate new genomic sequencing efforts.

The most important challenge for our study was the time required to retrieve the geographic coordinates associated with each Cytb sequence. Most sequences in open access databases such as GenBank do not have their latitude and longitude associated with them, which limits studies of spatial genetic variation. Efforts such as phylogatR (Pelletier et al. 2022) aggregate genetic sequences associated with georeferenced occurrences from global databases using automated pipelines, but researchers providing the sequences must include the proper metadata when uploading their data for this approach to work. We encourage scientists to share spatial data and museum catalogue numbers when applicable along with sequences when they are published, ensuring data reproducibility and extendibility.

### Conclusions

Our study underscores the value of integrating spatial information into macrogenetic studies. Our results support the hypotheses that variables associated with population size are positively correlated with genetic variation and that salamanders tend to exhibit spatial patterns of genetic variation that also differ across biogeographic realms. Based on our results, we reject the hypothesis that traits associated with reproductive strategies predict mitochondrial genetic variation in salamanders; and we partially support the hypothesis that older species tend to accumulate higher generic variation, as explained above. Our results suggest the importance of range-wide climatic variation, local environmental heterogeneity, and historical demographic events for influencing genetic variation. Future amphibian macrogenetic and genomic studies can incorporate more explicit tests of range expansion and contraction to better understand these processes. Finally, our findings have broad implications for establishing conservation priorities and highlighting regions where sampling has been historically scarce, particularly in biodiversity hotspots that warrant further attention.

## Data and code availability statement

The data and code (e.g., R scripts) that support the results of this research are openly available at https://github.com/luchoamador/amphibian_macrogenetics. Large folders and other supplementary materials are available in Dryad https://doi.org/10.5061/dryad.51c59zwjx.

## Notes

https://github.com/luchoamador/amphibian_macrogenetics

https://doi.org/10.5061/dryad.51c59zwjx

### Competing Interest Statement

The authors have declared no competing interest.

### Summary of Updates

The main changes incorporated in this version are: We include species age (divergence time) as a predictor of genetic variation, based on the Stewart and Wiens (2025) study (https://doi.org/10.1016/j.ympev.2024.108272). We discuss the geological events that could have influenced the genetic variation of the salamander species. We have included the statement –Data and Code Availability Statement-in the main text. We have carefully checked the alignments and species distribution ranges for all species and we revised the analyses based on the new findings. For example, our dataset increased from 214 species to 220 species. We modified the title following the new results. The new title is: Caudata macrogenetics: Geographic attributes and lineage age predict global patterns of mitochondrial genetic variation in salamanders.

### Funding

- National Science Foundation: DEB-2112946, DBI-1911293

